# *In vitro* and *in vivo* gene introduction in the cloudy catshark (*Scyliorhinus torazame*), a cartilaginous fish

**DOI:** 10.1101/2022.05.16.491766

**Authors:** Chika Fujimori, Chie Umatani, Misaki Chimura, Shigeho Ijiri, Hisanori Bando, Susumu Hyodo, Shinji Kanda

**Author notes:** Corresponding authors: Chika Fujimori, and Shinji Kanda.

## Abstract

Cartilaginous fishes have various unique physiological features such as cartilaginous skeletons and a urea-based osmoregulation strategy for adaptation to their marine environment. Also, because they are considered a sister group of bony vertebrates, understanding their unique features is important from an evolutionary perspective. However, experimental approaches are limited in cartilaginous fishes. Particularly, genetic engineering, which can analyze gene functions as well as cellular behavior, has not been effectively utilized in cartilaginous fishes. This is partly because their reproductive strategy involves internal fertilization, which results in difficulty in microinjection into fertilized eggs at the early developmental stage. Trials of gene transfer have also been limited both in *in vitro* cultured cells and *in vivo*. Here, to identify efficient gene transfer methods in cartilaginous fishes, we examined the effects of various methods both *in vitro* and *in vivo* using the cloudy catshark, a candidate model cartilaginous fish species. In all methods, green fluorescent protein (GFP) expression was used to evaluate exogenous gene introduction. First, we established a primary cell culture containing fibroblast-like and epithelial-like cells from cloudy catshark embryos. Using these primary cultured cells, we attempted gene transfection by lipofection, polyethylenimine (PEI), adenovirus, baculovirus and electroporation. Among the methods tested, lipofection, electroporation and baculovirus infection enabled the successful introduction of exogenous genes into primary cultured cells, allowing us to study physiological mechanisms at a single-cell level in culture conditions close to those in a living cartilaginous fish. We also attempted *in vivo* transfection into cloudy catshark embryos by electroporation and baculovirus infection. Although baculovirus-injected groups did not show GFP fluorescence, electroporation successfully introduced GFP into various tissues including muscle cells. Furthermore, we succeeded in GFP introduction into adult testis by electroporation. The *in vitro* and *in vivo* gene introduction methods that worked in this study may identify paths for future genetic manipulation including knockout experiments and cellular linage analysis in cartilaginous fishes.

## Introduction

Cartilaginous fishes have various unique biological characteristics such as cartilaginous skeletons and a urea-based osmoregulation strategy for adaptation to their marine environment. They also show diverse reproductive strategies, including oviparity and placental viviparity [1] [2] [3]. These fishes are considered as an extant sister group of bony vertebrates, which means that a thorough understanding of their biology will yield important information on the evolution of vertebrate physiology [4] [5] [6]. Despite the importance of cartilaginous fish research from this and other perspectives, our understanding of physiology of the cartilaginous fish remains relatively rudimentary. This may be due to the several characteristics of cartilaginous fish that make them difficult to study, including their long life cycle, their largely pelagic habitat and their large size.

Because these characteristics are minimized in the cloudy catshark (*Scyliorhinus torazame*), it is a promising candidate for a model cartilaginous fish species. First, they are rather small and easy to maintain in a laboratory aquarium. Second, they are oviparous and spawn approximately once every two to three weeks in laboratory tanks, such that developing embryos are easily obtained. Furthermore, genomic data acquired through whole-genome sequencing is available, which makes various molecular biological analyses easier [4].

Even in this model species, however, the limitations on experimental approaches remain a serious issue. In particular, genetic manipulation is difficult, so there have been no reports of the generation of transgenic or knockout cartilaginous fishes. The main factor limiting genetic manipulation of these species is their reproductive strategy: all cartilaginous fish species reported to date are copulating animals [1] [7]. They reproduce by internal fertilization; in oviparous species, the fertilized egg is encapsulated in an egg case and retained in the oviduct for a certain period before egg-laying occurs. In viviparous cartilaginous fishes, meanwhile, reproductive cycle including ovarian cycle and gestation period have been postulated several years [1] [7]. These reproductive features make it difficult to perform microinjection into fertilized eggs at early developmental stages. To overcome this obstacle, an important first step is establishing an efficient method of introducing exogenous genes into cultured cells and/or *in vivo* tissues. Thus far, however, there has been only one report of gene transfection into cartilaginous fish cells *in vitro*, in which the researchers attempted lipofection and electroporation to a cell line derived from dogfish shark embryos [8]. Furthermore, there is only one report of *in vivo* transfection in cartilaginous fishes using little skate embryos [9]. Thus, both *in vitro* and *in vivo* gene transfection are still challenging in cartilaginous fishes.

For exogenous gene introduction into vertebrate cells, three methods are commonly used: physical methods such as electroporation; chemical methods such as lipofection or the use of cationic polymers or polyethylenimine (PEI); and biological methods using viruses. Gene transfection using viruses has been reported in a limited number of species [10] [11] [12] and has never been attempted in cartilaginous fishes. In the present study, to identify efficient methods of gene transfer in cloudy catsharks, we first established a primary cell culture of cloudy catshark embryos and attempted gene transfection by lipofection, PEI, electroporation, adenovirus and baculovirus. We then attempted *in vivo* gene transfection into cloudy catshark embryos using the two methods that had been successful for *in vitro* transfection, namely, electroporation and baculovirus. Finally, *in vivo* gene transfection into adult testis was examined using electroporation.

## Materials and Methods

### Animals

Captive cloudy catsharks (*Scyliorhinus torazame*) and their naturally spawned eggs were transported from Ibaraki Prefectural Oarai Aquarium to the Atmosphere and Ocean Research Institute at the University of Tokyo. They were reared in a 1000 L tank with recirculating natural seawater at 16 °C under a constant photoperiod of 12 hours light/12 hours dark. The embryos were staged according to a method previously used for lesser spotted dogfish (*S. canicula*) [13] with the following slight modification: stage 31 was subdivided into two phases, stage 31 early (31E) and late (31L), as previously described [14]. All experiments were conducted in accordance with the Guidelines for Care and Use of Animals approved by the ethics committee of the University of Tokyo (P19-2).

### Primary cell culture

Embryos at stage 29-31E (before the pre-hatching period) were anesthetized with 0.02% ethyl 3-aminobenzoate methanesulfonate (MS-222) (Sigma-Aldrich, St. Louis, MO) in natural seawater and sacrificed by decapitation. After they had been cut into small pieces with sterilized scissors, embryonic tissues were incubated in a dissociation medium [L-15 medium (FUJIFILM Wako Pure Chemical Corporation, Osaka, Japan) containing 500 U/ml collagenase (FUJIFILM Wako Pure Chemical Corporation), 300 mM urea and 212.5 mM NaCl] for 1 hour at 16°C with pipetting every 15 minutes. Cell suspension was washed with culture medium and spread onto a 24-well culture plate coated with 0.1% gelatin. The culture medium containing L-15 supplemented with 20 mM HEPES (pH 7.4), 300mM urea, 180 mM NaCl, 20% FBS, 100 U/ml penicillin and 100 μg/ml streptomycin was replaced every three to four days. The osmolality of the dissociation medium and the culture medium was measured with a 5600 vapor pressure osmometer (Wescor, Logan, UT) and adjusted to ∼1000 mOsm with NaCl.

### Gene transfection to primary cultured cells

#### Lipofectamine and PEI transfection

Two days after primary cell culture was started, 0.8 μg of pEGFP-N1 vector (Takara Bio USA, Mountain View, CA), which carries EGFP under the promoter of human *Cytomegalovirus* (CMV), was transfected using 2 μl of Lipofectamine® 2000 (Thermo Fisher Scientific, Waltham, MA) or 2.5 μl of PEI-max (1.0 mg/ml) (Polysciences, Warrington, PA). Opti-MEM (Thermo Fisher Scientific) supplemented with 300 mM urea and 225 mM NaCl (∼1000 mOsm) was used as the medium for transfection. After seven days of transfection, cells were examined for GFP fluorescence.

#### Adenovirus infection

Recombinant adenovirus for the expression of EGFP was constructed according to the method described by Toyoda *et al*. [15] with minor modifications. In brief, CRE-nls-GFP ORF was amplified from addgene plasmid #49056 and inserted into the adenoviral expression vector pAd/CMV/V5-DEST^TM^ Gateway Vector plasmid (Thermo Fisher Scientific). After production and amplification of the adenovirus, the virus was purified using an Adenovirus (Ad5) Purification and Concentration Kit (AdenoPACK 20; Sartorius, Goettingen, Germany) according to the manufacturer’s instructions and stored at −80 °C until use. The titer of adenovirus was examined using an Adeno-XTM Rapid Titer Kit (Takara Bio USA) and determined to be 3.4 × 10^11^ ifu/ml. Two days after seeding the primary culture, 1 μl/well of the adenovirus was infected into the cells. Seven days after the transfection, cells were examined for GFP fluorescence.

#### Baculovirus infection

A bacmid containing inverted repeats of *piggyBac* transposon, CMVp-AcGFP and SV40p-Hyg (a hygromycin resistance gene) was constructed through modification of a pFastBac1-Δpolh plasmid [16]. These sequences were obtained from pPIGA3GFP (inverted repeats of *piggyBac* transposon [17]) and pAcGFP-Hyg C1 vector (CMVp-AcGFP and SV40p-Hyg) (632492; Takara Bio USA). Recombinant baculovirus was then produced using a Bac-to-Bac system. Briefly, the constructed plasmid was transformed into DH10Bac (Thermo Fisher Scientific). After transformation, recombinant bacmid DNA was purified using a Miniprep kit. The purified bacmid was confirmed through agarose gel electrophoresis. The recombinant bacmid was transfected into Sf9 cells using Cellfectin® II reagent (Thermo Fisher Scientific). The Sf9 cells were incubated for 120 hours at 28 to produce P1 baculoviruses, which were infected into Sf9 cells; these Sf9 cells were then incubated for 48 hours to yield P2 baculoviruses. After incubation, the supernatants of the P2 incubation media were collected and used for the following infection experiments. The titer of baculovirus was measured by GFP expression on HEK293A cells and 1.55 × 10^5^ gene transfer unit (gtu)/ml of P2 supernatant was obtained. In the experiment validating this transfection method (Fig. 2), 5 μl of P2 supernatant was treated. To examine the infection efficiency of baculovirus (Fig. 3),2-20 μl of P2 supernatants were added to cloudy catshark primary cultured cells two days after the culture was started. We examined the cells for GFP fluorescence seven days after infection.

**Fig 1.**
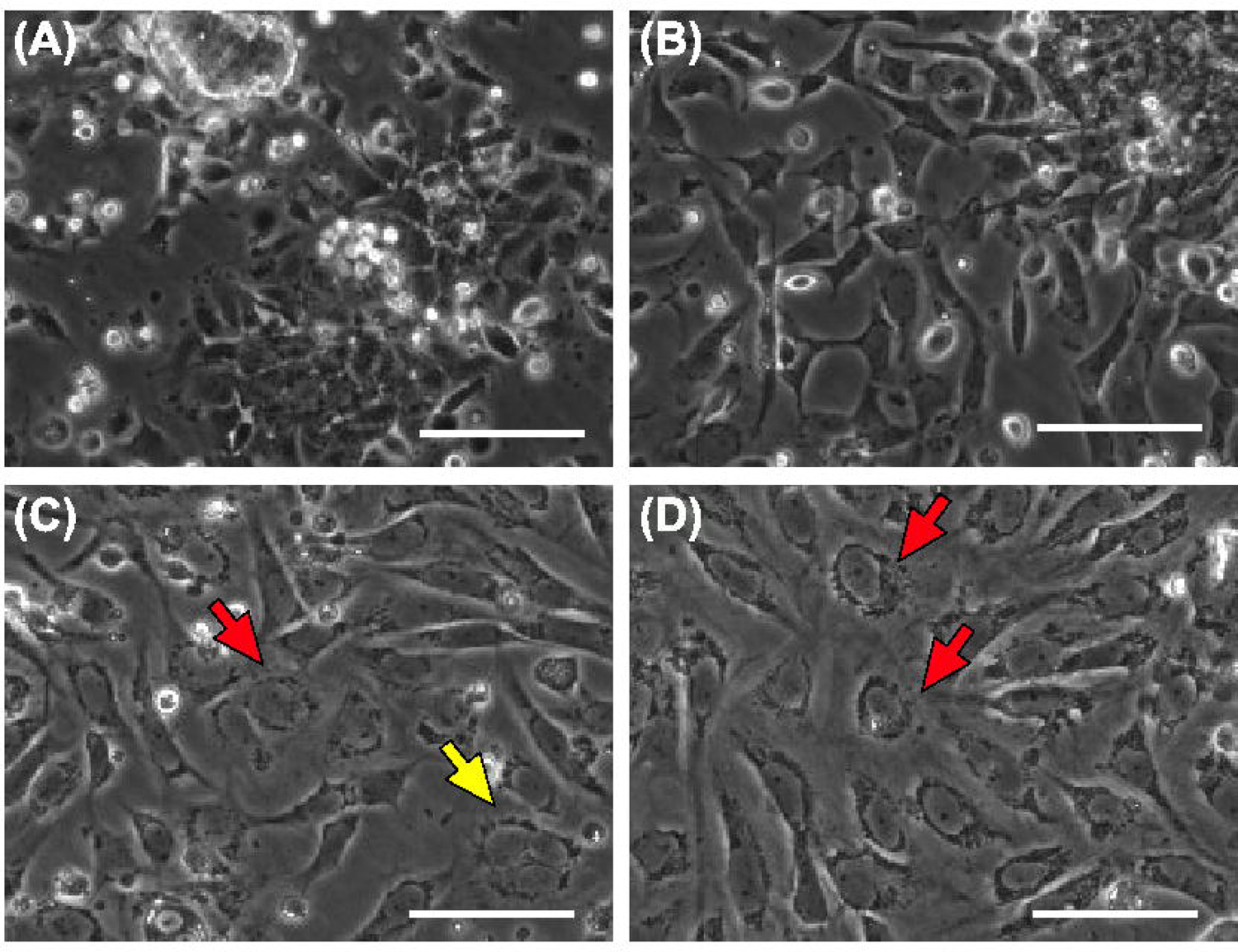
Primary cultured cells from cloudy catshark embryos. Morphological observation of the primary cultured cells from cloudy catshark embryos at 1 (**A**), 9 (**B**), 19 (**C**) and 28 (**D**) days after start of culture. **A, B** Elongated fibroblast-like and polygonal epithelial-like cells had spread from cell clusters into a monolayer. **C, D** After 19 days of culture, multinucleated cells (yellow arrow), flattened nuclei and cytoplasms, and vacuoles around the nuclei were observed (red arrows). The cells could be cultured for at least 28 days under the same culture conditions. Scale bars represent 100 μm.

**Fig 2.**
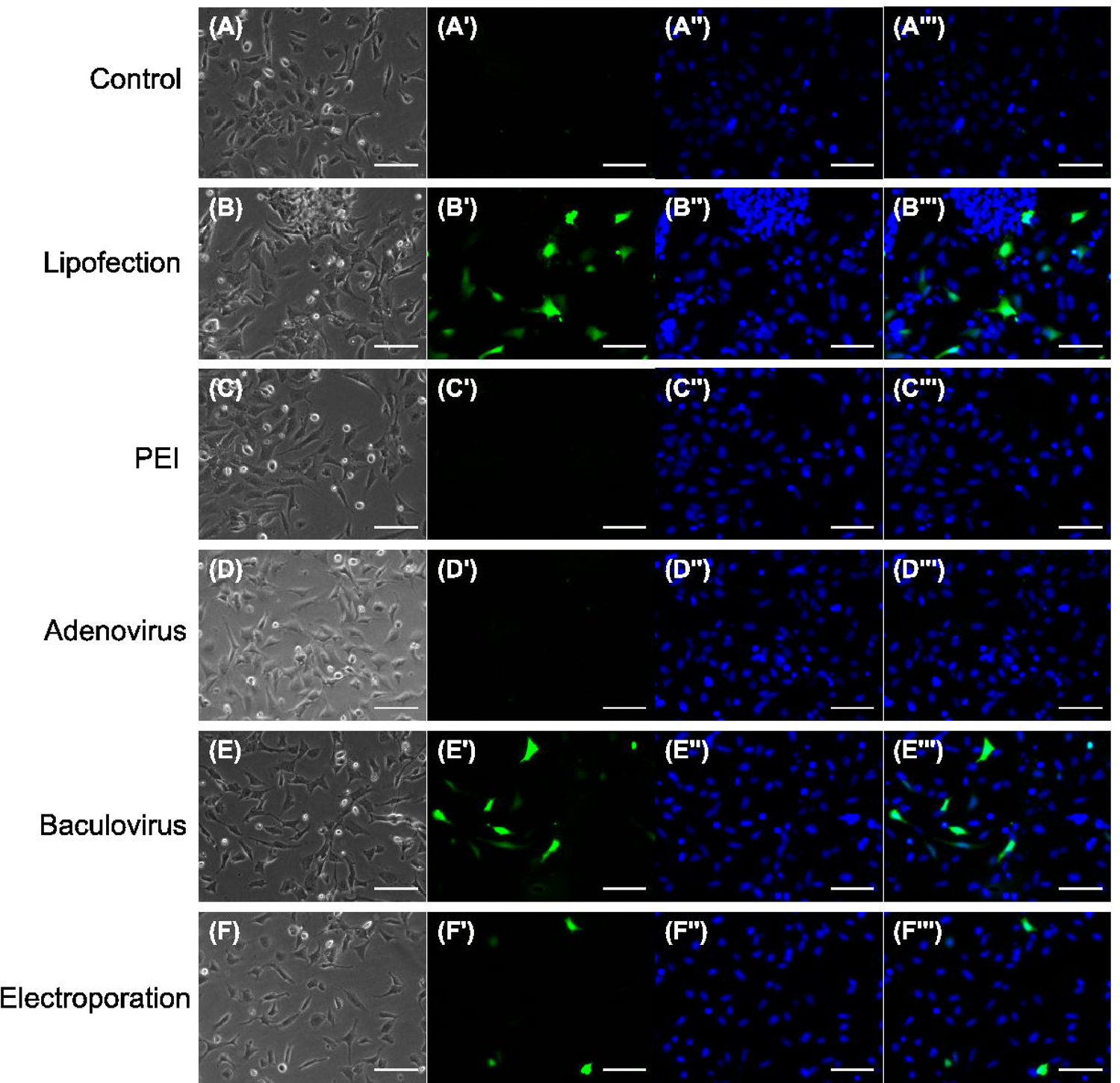
Gene transfection to primary cultured cells of cloudy catshark by various methods. The morphology of primary cultured cells after transfection (**A-F**) and their GFP fluorescence images (**A’-F’**). Their nuclei were counterstained with Hoechst 33342 (**A’’-F’’**) and rightmost columns show the merged images with GFP fluorescence (**A’’’-F’’’**). Primary cultured cells were transfected by lipofection (**B-B’’’**), PEI (**C-C’’’**), adenovirus (**D-D’’’**), baculovirus (**E-E’’’**) or electroporation (**F-F’’’**). GFP-expressing cells were observed among the cells transfected by lipofection (**B’**), baculovirus infection (**E’**) and electroporation (**F’**). **A** was untransfected cells. Scale bars represent 100 μm.

**Fig 3.**
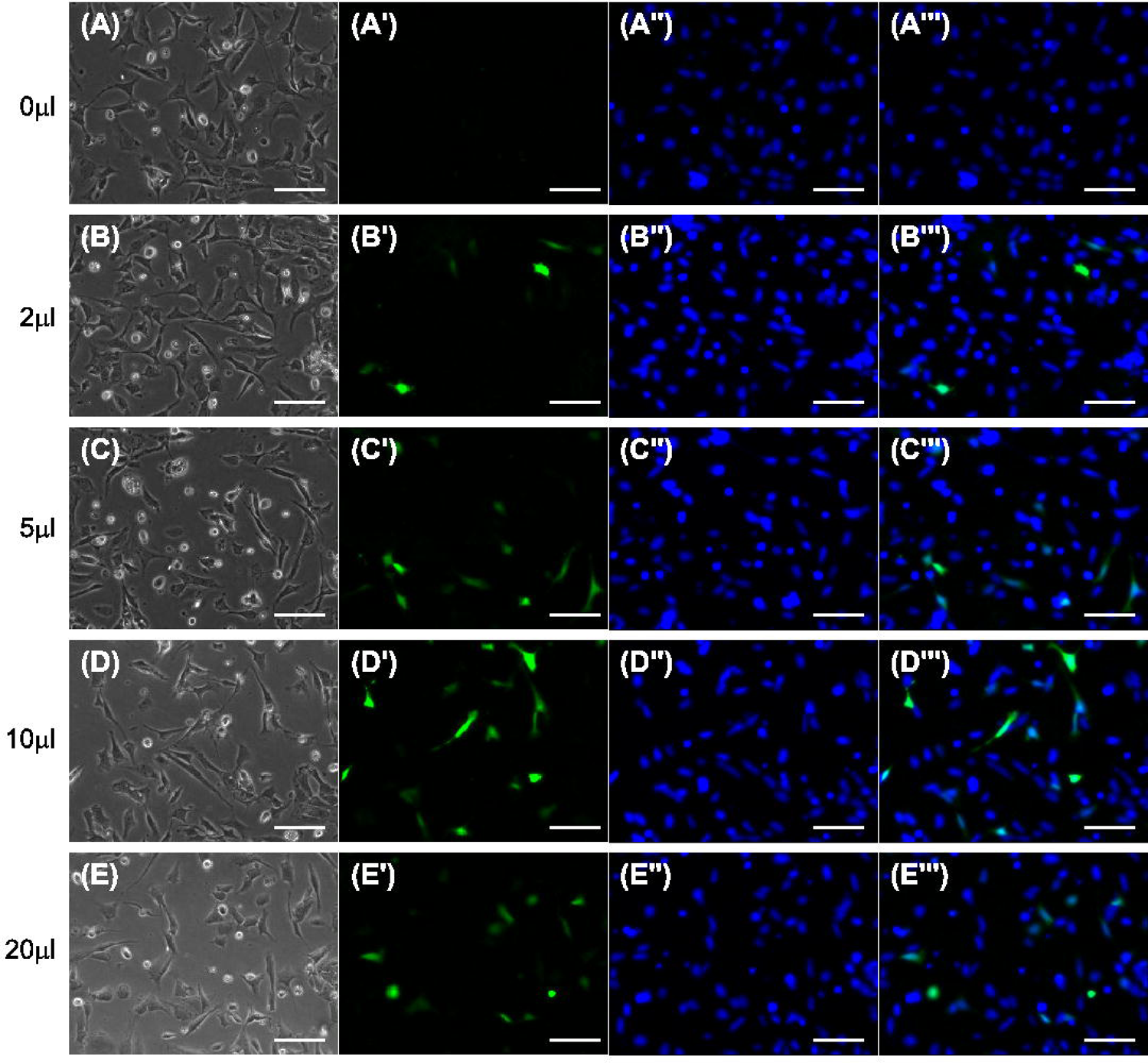
Dose-dependent increase in efficiency of baculovirus-mediated GFP transfection *in vitro*. Phase-contrast images (**A-E**) and the corresponding GFP fluorescent images (**A’-E’**) seven days after infection with various quantities of baculovirus. Hoechst 33342 was used for nuclear counterstaining (**A’’-E’’)** and merged images with GFP fluorescence were shown in **A’’’-E’’’.** The volume of baculovirus solution: **A-A’’’,** 0 μl; **B-B’’’**, 2 μl; **C-C’’’**, 5 μl; **D-D’’’**, 10 μl; **E-E’’’**, 20 μl. The titer of the P2 supernatant of baculovirus on HEK293A cells was 1.55 × 10^5^ gtu/ml. Scale bars represent 100 μm.

### Electroporation

Electroporation was performed immediately after cell dissociation. A cell suspension containing 50 ng/μl of pEGFP-N1 vector was inserted into 1-mm gap cuvettes and chilled on ice for 10 minutes. Electroporation was then conducted under various conditions using a T820 electroporator (BTX, Holliston, MA). The cell suspension was then chilled on ice for another 10 minutes and seeded onto 0.1% gelatin-coated dishes. In the experiment validating this transfection method (Fig. 2), electroporation was performed at 0.2 kV for 50 μsec with three pulses. We examined the cells for GFP fluorescence nine days after electroporation.

### Examination of transfection efficiency in cell culture

Nine days after the start of cell culture, Hoechst 33342 (NucBlue™ Live ReadyProbes™ Reagent, Thermo Fisher Scientific) was added according to the manufacturer’s instructions. Then, 21.4 mm^2^ of the contents of each well was photographed through an All-in-One Fluorescence Microscope BZ-X810 (KEYENCE, Osaka, Japan) equipped with the following filter set: Hoechst, ex. 360 ± 20/ em. 460 ± 25; GFP (EGFP, AcGFP), ex. 470 ± 20/ em. 525 ± 25. The numbers of total cells and fluorescence-positive cells within each captured area were counted using ImageJ. Note that a very small percentage of cells showed autofluorescence. These inevitable pseudo-positive cells can be estimated from the control samples without transfection.

### In vivo transfection to cloudy catshark embryos and adult testis

Embryos at stage 31L to 32 were anesthetized with 0.02% MS-222 in natural seawater, and baculovirus infection or electroporation was performed. One to two μl of the P2 suspension, which is the same suspension used for the primary cultured cells (for baculovirus infection), or 0.5-2 μg of pEGFP-N1 vector (for electroporation) was injected into a subcutaneous site using a 10 μl Hamilton syringe #80300 (Hamilton Company, Reno, NV). For electroporation, embryos were temporarily transferred to PBS, which is in lower ionic conditions than their body fluid, to reduce leakage of the given charge out of the embryos. Electroporation was carried out using a handmade tweezer-type electrode set with 10 mm × 8 mm aluminum plates and a T820 electroporator at 15-50 V for 50 ms with two to eight pulses. After transfection, all embryos were immediately reared in seawater at 16 °C with aeration for seven days; seven days after gene introduction, we examined the embryos for GFP fluorescence. For *in vivo* electroporation to adult testis, histological analysis of the cloudy catshark testis indicated that many undifferentiated germ cells are localized in the germinal zone running longitudinally through the testis (Additional file 3A and B, arrows), which is consistent with previous reports in other cartilaginous fishes [1] [18] [19]. Adult catshark (267.5 g–454.3 g) were anesthetized with 0.02% MS-222 in natural seawater, and one of the testes was exposed from a small incision (∼5 cm) in the abdomen. Then, 50-100 μg of pEGFP-N1 vector was injected into the germinal zone in the dorsal testis (Additional file 3A, arrows), and 5 electric pulses were applied at 30 V for 50 ms pulse with the electrode used in embryo *in vivo* electroporation. After the electroporation, the incision was sutured, and they were reared in 16 °C seawater with aeration. After 5-7 days, the cloudy catsharks were anesthetized and sacrificed. Their electroporated tissues were sampled, and GFP fluorescence was observed.

### Immunohistochemistry

Five to seven days after *in vivo* transfection, cloudy catshark embryos or adults were anesthetized with 0.02% MS-222. The injected areas were dissected out and fixed with 4% paraformaldehyde in 0.05 M PBS. After cryosectioning [20], immunohistochemistry was performed according to a slightly modified version of the protocol described by Takahashi *et al.* [21]. Briefly, 25 (embryos) or 30 (adults) μm-thick transverse cryosections were rinsed with PBS containing 0.5% triton X-100 (PBST) and incubated with anti-GFP rat monoclonal antibodies (1:1000, GF090R; Nacalai Tesque Inc., Kyoto, Japan). After endogenous peroxidase activity was inactivated by the addition of 0.3% H_2_O_2_ in PBS, a secondary antibody reaction was performed using anti-rat IgG biotinylated antibodies (1:200, Jackson ImmunoResearch, West Grove, PA). Next, sections were incubated with VECTASTAIN® Elite ABC-HRP kit reagent (Vector Laboratories). GFP expression cells were visualized with 3,3-diaminobenzidine (DAB) and 0.003% H_2_O_2_, or Alexa Fluor® 555 Streptavidin (Thermo Fisher Scientific). The sections of the embryo brain were counterstained with cresyl violet acetate after DAB reaction. For adult testes, transverse sections were prepared, and they were stained with hematoxylin and eosin as previously described [14]. The stained sections were photographed with an All-in-One Fluorescence Microscope BZ-X810 (KEYENCE, Osaka, Japan) with its automatic image stitching function.

### Statistical analysis

All values are shown as mean ± standard deviation (SD). The transfection efficiencies of baculovirus infection and electroporation in cloudy catshark primary cell culture were analyzed using Dunnett’s multiple comparison test, and the significance of the difference from control conditions was assessed. *p*-values less than 0.05 were considered statistically significant.

## Results

### Primary cell culture from cloudy catshark embryos

We used embryos at stage 29-31E for the primary cell culture to avoid pre-hatching, which occurs at the end of stage 31E [13] and may increase the risk of microbial contamination due to the influx of seawater to the inside of the egg capsule. Using dissociated cells from these embryos, we successfully established a primary culture in a standard L-15 medium supplemented with FBS, urea and NaCl. Although the cells were initially scattered in many clusters, they spread out from their clusters into a monolayer within 24 hours of the culture was started (Fig. 1A and Additional file 1). The cultured cells were heterogeneous, and the most abundant types were elongated fibroblast-like cells and polygonal epithelial-like cells (Fig. 1B). After 19 days of culture, the cultured cells became unhealthy. Specifically, the cytoplasms and nuclei of some cells became enlarged and flattened, and vacuoles were observed around the nucleus (Fig. 1C, red arrows). In addition, multinucleated cells were observed (Fig. 1C, yellow arrow). These characteristics were similar to those of senescent cells [22]. The cells could be cultured for at least 28 days under the same culture conditions, although the proportion of cells considered senescent increased over time (Fig. 1D).

### Exogenous gene transfection into primary cultured cells from cloudy catshark embryos

We used the primary cells established in this study to examine various methods of gene transfection. All constructs and viruses incorporated the ubiquitous promoter CMVp and EGFP/AcGFP, and we assessed the efficiencies of the various methods by examining the quantity of GFP expression in each batch of transfected cells under a fluorescent microscope. Primary cell cultures of cloudy catshark embryos contained very small numbers of cells showing autofluorescence. Since it was difficult to distinguish uncharacterized autofluorescence from GFP signals, we counted the total numbers of fluorescence-positive cells including both GFP-fluorescent and autofluorescent cells (Table 1). As Table 1 shows, only a very tiny proportion (0.018%) of total cells in the untreated group were fluorescence-positive, indicating autofluorescence (Fig. 2A-A’’’). Since the proportions of fluorescence-positive cells were much higher in the lipofection (4.669%)-, baculovirus (1.983%)- and electroporation (0.694%)-treated groups, we consider that most of the fluorescence-positive cells in these groups can be assumed to be EGFP/AcGFP-expressing cells (Table 1). In the lipofection-treated group using lipofectamine® 2000, the first fluorescent cells were seen two days after transfection; on day seven after transfection, fluorescence was observed in about 4.669% of the cells (Fig. 2B-B’’’). When 0.8 μg plasmids were transfected with PEI, no GFP-expressing cells were observed (Fig. 2C-C’’’). Similarly, GFP-expressing cells were scarcely observed among the adenovirus-infected cells (Fig. 2D-D’’’). In baculovirus-infected cells, on the other hand, the first GFP-expressing cells appeared four days after infection; seven days after infection, 1.983% of the cultured cells expressed GFP (Fig. 2E-E’’’). Electroporation-treated cells also showed GFP signals from five days after electroporation, most abundantly on day 9, when 0.694% of the cultured cells were fluorescent (Fig. 2F-F’’’).

**Table 1.** The proportions of cells with fluorescence after GFP-containing vector or virus transfection by various methods.

|  | (number of fluorescent cells/all cells) | % of fluorescent cells | n |
| --- | --- | --- | --- |
| Control | (0/5295) | $0.018 \pm 0.021$ | 4 |
|  | (2/5866) |  |  |
|  | (2/5152) |  |  |
|  | (0/5443) |  |  |
| Lipofection | (178/4688) | $4.669 \pm 1.544$ | 4 |
|  | (231/3474) |  |  |
|  | (149/2934) |  |  |
|  | (104/3300) |  |  |
| PEI | (0/5377) | $0.005 \pm 0.009$ | 4 |
|  | (0/6732) |  |  |
|  | (1/5519) |  |  |
|  | (0/4826) |  |  |
| Adenovirus | (7/5474) | $0.036 \pm 0.062$ | 4 |
|  | (0/6083) |  |  |
|  | (0/5511) |  |  |
|  | (1/5879) |  |  |
| Baculovirus | (111/5075) | $1.983 \pm 0.560$ | 4 |
|  | (157/5886) |  |  |
|  | (83/5902) |  |  |
|  | (94/5625) |  |  |
| Electroporation | (54/4960) | $0.694 \pm 0.357$ | 3 |
|  | (26/4328) |  |  |
|  | (21/5344) |  |  |

Two of the gene-introducing methods that are successful in *in vitro* culture, baculovirus infection and electroporation, have previously been applied to *in vivo* gene transfection in mammals and teleost fish [23] [24] [25] [26]. We further tested the utility of these methods under various conditions using primary cell culture. Cell cultures that were infected with different amounts of baculovirus showed higher proportions of fluorescent cells in a dose-dependent manner: the proportions of cells showing GFP fluorescence after being treated with 0, 2, 5, 10 and 20 μl of baculovirus were 0.034% (Fig. 3A-A’’’; only autofluorescence), 0.685% (Fig. 3B-B’’’), 1.664% (Fig. 3C-C’’’), 2.882% (Fig. 3D-D’’’) and 3.576% (Fig. 3E-E’’’), respectively (Table 2). For the electroporation method, we tested various numbers, durations, and voltages of the pulses. The proportions of GFP-expressing cells after electroporation were as follows: 0.022% (0 kV, 0 μsec, 0 pulse; Fig. 4A-A’’’, only autofluorescence), 0.003% (0.05 kV, 50 μsec, 3 pulses; Fig. 4B-B’’’), 0.069% (0.1 kV, 50 μsec, 3 pulses; Fig. 4C-C’’’), 0.376% (0.2 kV, 50 μsec, 3 pulses; Fig. 4D-D’’’), 0.137% (0.2 kV, 10 μsec, 3 pulses; Fig. 4E-E’’’), 0.264% (0.2 kV, 25 μsec, 3 pulses; Fig. 4F-F’’’), 0.190% (0.2 kV, 50 μsec, 1 pulse; Fig. 4G-G’’’) and 0.383% (0.2 kV, 50 μsec, 2 pulses; Fig. 4H-H’’’). Cell cultures that were electroporated at 0.2 kV for 50 μsec with two pulses or at 0.2 kV for 50 μsec with three pulses had significantly higher proportions of fluorescence-positive cells compared to the control cultures, in which all fluorescence was autofluorescence (Table 3). Note that three pulses at 0.2 kV for 50 μsec appeared to cause severe damage to the cells, as many cells came unstuck after electroporation.

**Fig 4.**
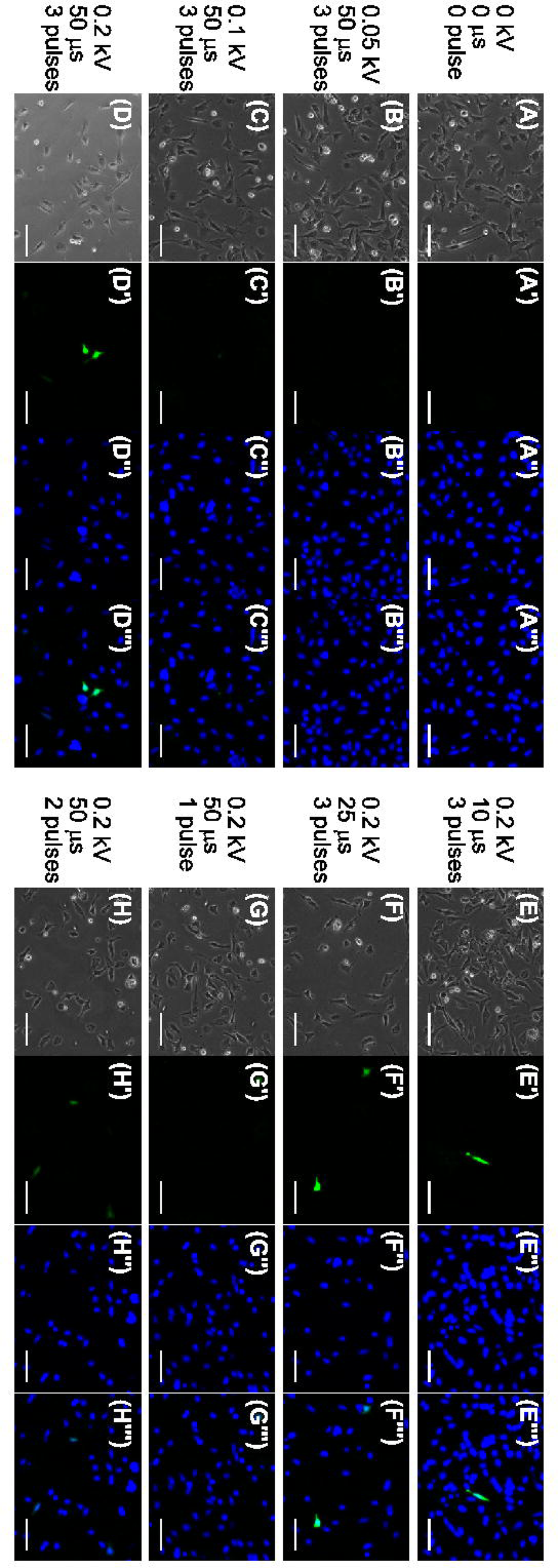
Efficiency of gene transfection under various conditions of electroporation in cloudy catshark cells *in vitro*. Phase-contrast images of cell morphology (**A-H**) and the corresponding fluorescence images (**A’-H’**) nine days after electroporation. Nuclear counterstaining images using Hoechst 33342 and their merged images with GFP fluorescence were shown in **A’’-H’’** and **A’’’-H’’’,** respectively. Electroporation was performed using different voltages (**A-A’’’**, 0 kV; **B-B’’’**, 0.05 kV; **C-C’’’**, 0.1 kV; **D-D’’’**, 0.2 kV), durations (**A-A’’’**, 0 μs; **E-E’’’**, 10 μs; **F-F’’’**, 25 μs; **D-D’’’**; 50 μs) and pulse numbers (**A-A’’’**, 0 pulse; **G-G’’’**, 1 pulse; **H-H’’’**, 2 pulses; **D-D’’’**, 3 pulses). When electroporation was performed at 0.2 kV, 50 μsec, 3 pulses (**D-D’’’**) and 0.2 kV, 50 μsec, 2 pulses (**H-H’’’**), the percentages of GFP-expressing cells were significantly higher than they were in the control (**A-A’’’**). Scale bars represent 100 μm.

**Table 2.** The proportions of fluorescent-positive cells after infection with different volumes of baculovirus. Statistically significant difference from control (cells treated with 0 μl baculovirus) was assessed by Dunnett’s multiple comparison test (\*\**p* < 0.01).

| Baculovirus (μl) | (number of fluorescent cells/all cells) | % of fluorescent cells | n |
| --- | --- | --- | --- |
| control (0 μl) | (7/6690) | $0.034 \pm 0.048$ | 4 |
|  | (1/6968) |  |  |
|  | (0/5960) |  |  |
|  | (1/5421) |  |  |
| 2 | (47/5910) | $0.685 \pm 0.201$ | 4 |
|  | (58/6591) |  |  |
|  | (22/5208) |  |  |
|  | (41/6390) |  |  |
| 5 | (103/6200) | $1.664 \pm 0.601$ | 4 |
|  | (139/5880) |  |  |
|  | (48/5354) |  |  |
|  | (108/6223) |  |  |
| 10 | (154/6239) | $2.882 \pm 1.444^{**}$ | 4 |
|  | (246/4935) |  |  |
|  | (127/5330) |  |  |
|  | (102/6026) |  |  |
| 20 | (171/6058) | $3.576 \pm 1.810^{**}$ | 4 |
|  | (352/5599) |  |  |
|  | (80/3105) |  |  |
|  | (139/5310) |  |  |

**Table 3.** The proportions of cells with fluorescence after electroporation under various conditions. Statistically significant difference from control (cells subjected to 0 kV electroporation) was assessed by Dunnett’s multiple comparison test (\*\**p* < 0.01).

| Voltage (kV) | Length (μs) | Number | (number of fluorescent cells/all cells) | % of fluorescent cells | n |
| --- | --- | --- | --- | --- | --- |
| 0 | 0 | 0 | (6/8362) | 0.022 ± 0.034 | 4 |
|  |  |  | (1/6224) |  |  |
|  |  |  | (0/6397) |  |  |
|  |  |  | (0/7587) |  |  |
| 0.05 | 50 | 3 | (1/8858) | 0.003 ± 0.006 | 4 |
|  |  |  | (0/5882) |  |  |
|  |  |  | (0/5774) |  |  |
|  |  |  | (0/7760) |  |  |
| 0.1 | 50 | 3 | (7/9491) | 0.069 ± 0.028 | 4 |
|  |  |  | (5/6516) |  |  |
|  |  |  | (5/5147) |  |  |
|  |  |  | (2/6780) |  |  |
| 0.2 | 50 | 3 | (4/8468) | 0.376 ± 0.272** | 4 |
|  |  |  | (32/4507) |  |  |
|  |  |  | (17/4215) |  |  |
|  |  |  | (12/3499) |  |  |
| 0.2 | 10 | 3 | (7/8261) | 0.137 ± 0.121 | 4 |
|  |  |  | (20/6290) |  |  |
|  |  |  | (4/6207) |  |  |
|  |  |  | (6/7552) |  |  |
| 0.2 | 25 | 3 | (9/6882) | 0.264 ± 0.144 | 4 |
|  |  |  | (21/4547) |  |  |
|  |  |  | (10/3699) |  |  |
|  |  |  | (14/7213) |  |  |
| 0.2 | 50 | 1 | (11/8196) | 0.190 ± 0.046 | 4 |
|  |  |  | (14/5788) |  |  |
|  |  |  | (9/4383) |  |  |
|  |  |  | (12/6793) |  |  |
| 0.2 | 50 | 2 | (22/6628) | 0.383 ± 0.208** | 4 |
|  |  |  | (31/4577) |  |  |
|  |  |  | (11/5846) |  |  |
|  |  |  | (17/5084) |  |  |

### In vivo transfection into cloudy catshark embryos

Following our methodological validation using primary cultured cells, we next attempted *in vivo* gene introduction to embryos at stage 31L-32 through baculovirus infection and electroporation. We chose these stages for *in vivo* transfection because they are sufficiently grown-up to exhibit many physiological processes such as nutritional absorption through the digestive tract, buccal pumping, and a functional immune system, yet their skin is still not too thick [14] [27] [28]. After seven days of electroporation, GFP fluorescence was clearly observed in the skeletal muscles of the injected areas (Fig. 5A and B). A dot-like pattern of fluorescence on the skin surfaces was observed in both untreated (Fig. 5H) and electroporated embryos (Fig. 5B), indicating that this fluorescence is autofluorescence. Immunohistochemistry using EGFP antibodies confirmed that EGFP was expressed in some muscular cells in the vector-injected side (Fig. 5C), while not expressed in those in the contralateral side (Fig. 5D). In addition to muscular cells, EGFP was also detected in neuron-like cells in the embryos that we performed electroporation in the brain. After sectioning, neurons in the subpallium of the brain were found to be labeled by GFP (Additional file 2). In embryos infected with baculovirus, on the other hand, no obvious GFP signals were observed at the injection sites (Fig. 5E, F and G), similar to the situation in uninjected embryos (Fig. 5H). Next, we examined the survival and GFP fluorescence one week after electroporation with various plasmid volumes and numbers and voltages of the pulses. When electroporation was performed using 0.5 μg (Fig. 6A), 1 μg (Fig. 6B) or 2 μg vector (Fig. 6C), GFP fluorescence was observed in all conditions, and there was no apparent difference in the transfection efficiency. The embryos pulsed at 15 V were viable, and GFP fluorescence was observed in both 5 pulsed and 8 pulsed conditions. For embryos pulsed at 30 V, one of two embryos treated with 8 pulses survived and showed GFP fluorescence, and the other died within one week. When the embryos were electroporated at 30 V and 5 pulses, both of the two embryos remained alive and GFP fluorescence was observed. Under the 50 V pulse condition, both 2 and 5 pulses treated embryos died within one week.

**Fig 5.**
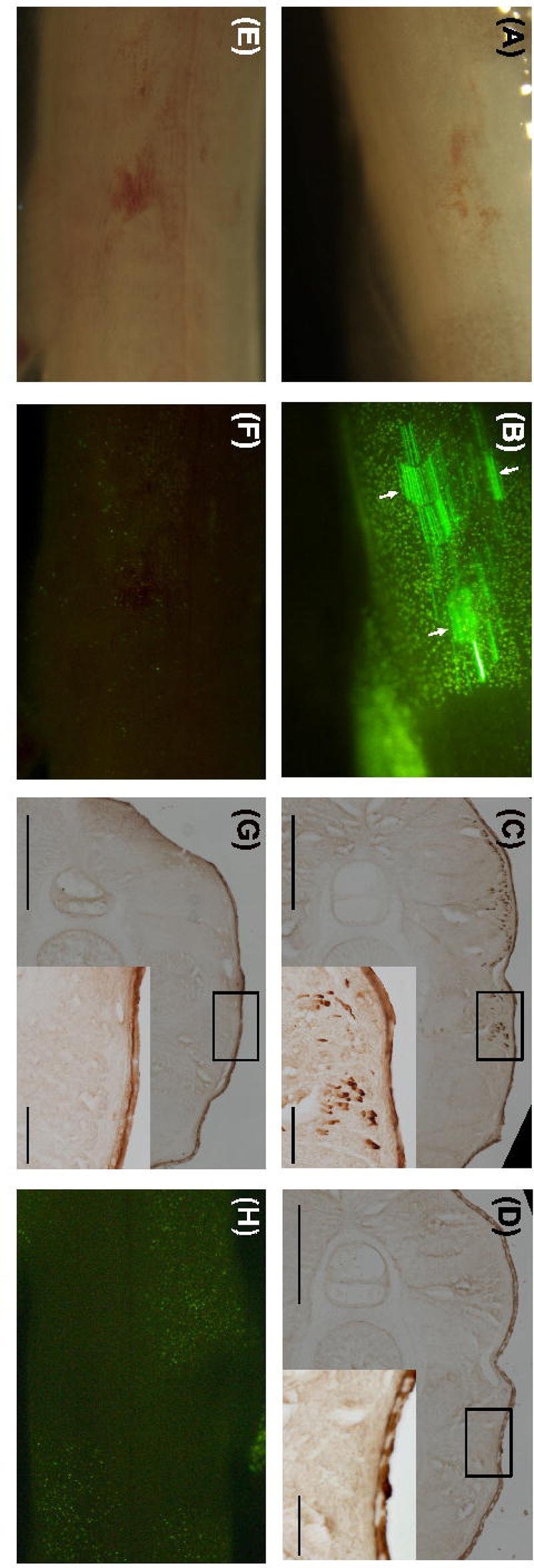
*In vivo* gene introduction into cloudy catshark embryos by electroporation or baculovirus infection. Bright field (**A, E**) and fluorescence images (**B, F**) of cloudy catshark embryos seven days after electroporation (**A, B**) or baculovirus infection (**E, F**). GFP fluorescence was observed in the muscle of the injected area of an embryo subjected to electroporation (**B**, arrows), but not in the specimen injected with baculovirus (**F**), or in untreated embryo (**H**). Note that dot fluorescence observed in (**B, F, H**) represents autofluorescence of skin. **C, D, G** Immunohistochemistry for EGFP of the transverse sections of embryos (left ventral to right dorsal images) also showed that electroporated (**C**), but not baculovirus-infected (**G**), specimens express EGFP. Insets show magnified images of boxed areas. EGFP-immunoreactive cells were only observed in plasmid injected-side of the electroporated embryos and were not observed in the contralateral side of the body (**D**). Note that **D** and **G** show lateral view of the cylinder-shaped muscle cells whereas **C, D** and **G** represent the transverse section of them. Scale bars represent 500 μm (**C, D, G**) and 50 μm (insets).

**Fig 6.**
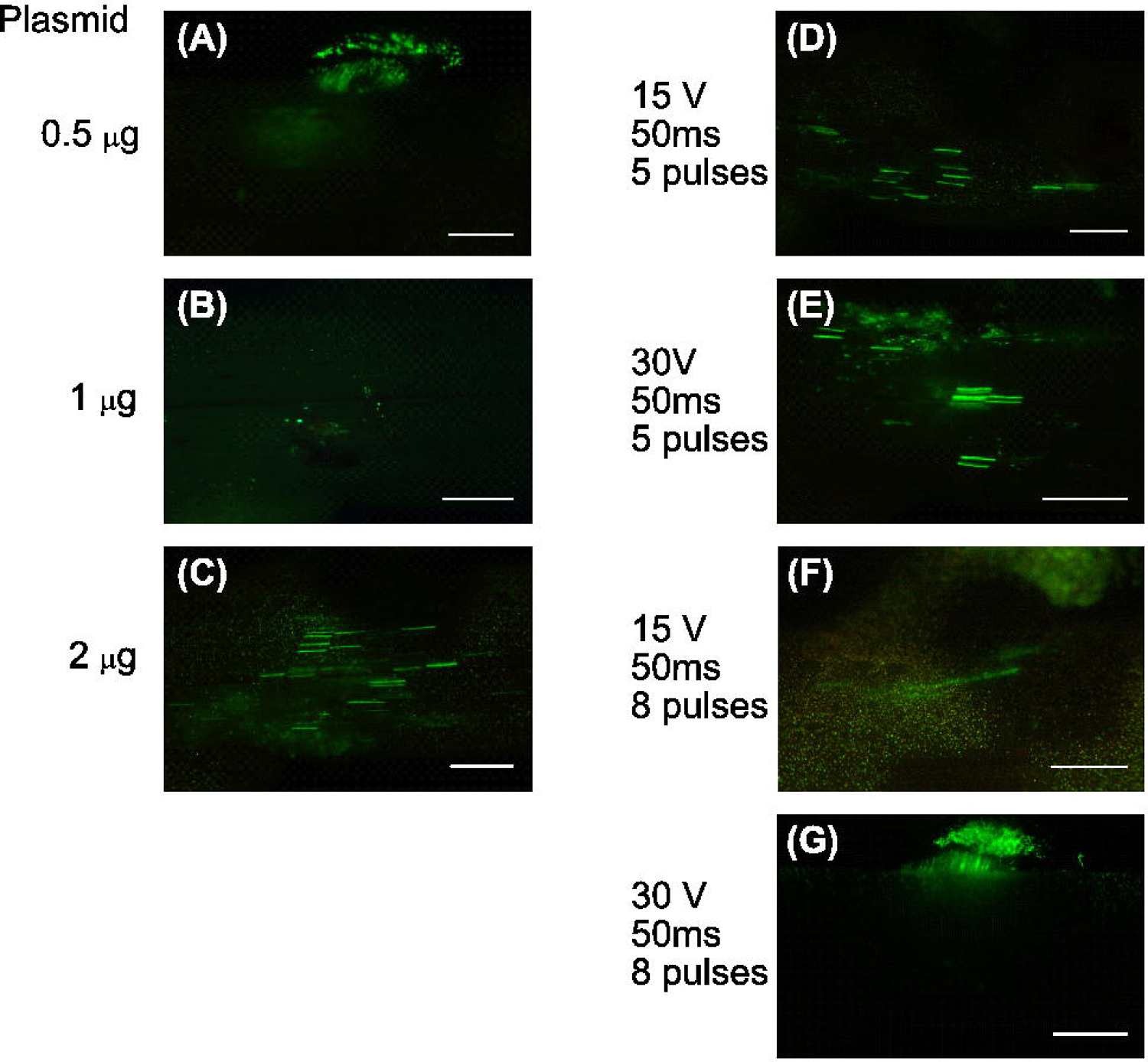
The effect on survival and gene transduction under various conditions of electroporation in cloudy catshark embryos *in vivo*. Fluorescence images of cloudy catshark embryos seven days after electroporation. Electroporation was performed using different plasmid volume (**A, D, E, F, G**, 0.5 μg; **B**, 1 μg; **C**, 2 μg), voltage **(D, F,** 15 V; **E, G,** 30 V), and pulse numbers (**D, E,** 5 pulses; **F, G,** 8 pulses). Note that the electroporated embryos at 50 V, 50 ms, 2 pulses (two of two embryos), 50 V, 50 ms, 5 pulses (two of two embryos), 30 V, 50 ms, 8 pulses (one of two embryos) died within seven days of electroporation. Scale bars represent 1 mm.

### In vivo transfection into adult cloudy catsharks

Finally, we attempted gene introduction in adult testis by electroporation at 30 V and 5 pulses. Based on the histological observation, we tried electroporation in the germinal zone where undifferentiated germ cells are localized (Additional file 3) [1] [18] [19]. One week after electroporation, GFP fluorescence was observed in testis (Fig. 7A), while GFP fluorescence was not detected in untreated testis (Fig. 7B). Cryosection of the electroporated testis indicated the GFP florescence was restricted outside the spermatocysts (Fig. 7C-E). In addition to the raw GFP signal (Fig. 7C, green), immunohistochemistry using GFP antibody (Fig.7D, magenta) showed red fluorescence in the same cell (magenta signal in Fig. 7E), which indicates this fluorescence is derived from GFP.

**Fig 7.**
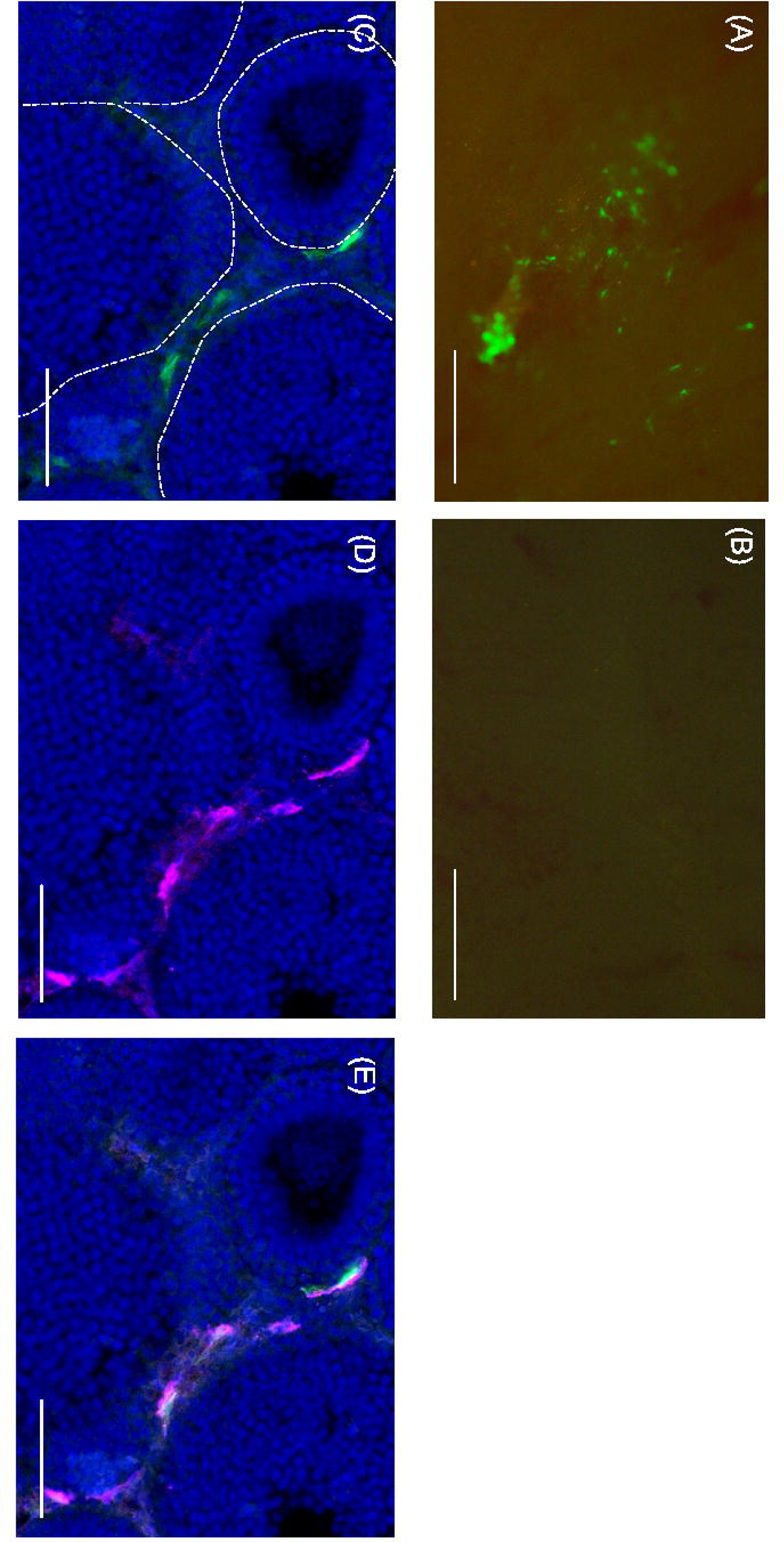
*In vivo* gene introduction into adult cloudy catshark testis. Fluorescence image of adult cloudy catshark testis seven days after electroporation (**A**) and untreated testis (**B**). **C, D, E** EGFP signals in the testis section. Raw GFP fluorescence (**C**), which is confirmed to show GFP-immunoreactivity (**D**: GFP**-**immunoreactive signal [magenta]**, E**: merge) outside of spermatocysts. Dashed lines indicate the sperematocysts. Scale bars represent 1 mm (**A, B**) and 200 μm (**C, D, E**), respectively.

## Discussion

In the present study, we generated primary cell cultures from cloudy catshark embryos and examined the effectiveness of various gene transfection methods *in vitro* and *in vivo*. We found that lipofection, baculovirus infection and electroporation can be used to introduce exogenous genes into primary cultured cells of the cloudy catshark. Electroporation can also be used for *in vivo* gene transfection into not only embryos but also adult testis; this finding represents for the first time that *in vivo* gene transfection has been successfully performed in adult cartilaginous fishes. The established primary culture system in combination with these newly verified *in vitro* and *in vivo* transfection methods will provide a novel tool for future investigations using genetic engineering at the cellular, tissue and individual levels in cartilaginous fishes.

### Cellular morphologies and properties of the primary cultured cell system

The vast majority of primary cultured cells from cloudy catshark embryos showed elongated fibroblast-like and polygonal epithelial cell-like morphology. Although it was difficult to identify the types of heterogeneous cellular populations in the primary culture, the observed morphologies were similar to those in embryonic primary cultures achieved in other animals, which mainly consist of epithelial and fibroblast cells [29] [30].

On the other hand, our time-lapse recordings rarely showed cell division, which is usually observed in primary cultured cells from embryos (Additional file 1). Although further quantitative examination using BrdU is necessary to confirm this, the frequency of cell division might be lower in our experiments than it has been in reports using other vertebrates including sharks [31] [32]. One possible reason for this low frequency of cell division is the low incubation temperature. Because the cloudy catshark specimens and their primary cultured cells were kept and incubated at 16 °C in this study, their metabolic and mitotic rates are expected to be low. In fact, the development of the cloudy catshark embryo takes a relatively long time: it takes six months to go from fertilization to hatching, during which time they only grow to ∼3.5 g [14] [33]. After 19 days of culture, we observed cells with the typical characteristics of senescence such as multinucleation, flattened nuclei and cytoplasms, and/or vacuoles around the nuclei [22]. This may be due to the lack of molecules that are originally present in the cartilaginous fish body fluid such as Trimethylamine-N-oxide (TMAO), or growth factors such as insulin and EGF. Future experiments with different culture conditions may improve the quality of the primary culture.

### Lipofection, baculovirus infection and electroporation were effective for gene transfection into cloudy catshark *cells*

Of the various transfection methods examined in this study, only lipofection, baculovirus infection and electroporation effectively introduced exogenous genes into cloudy catshark primary culture cells. This is the first report of successful viral transfection into cartilaginous fish cells. Baculovirus was originally discovered as a virus that infects insect cells and was not initially believed to infect vertebrate cells. Nevertheless, successful gene delivery by baculovirus vectors has been reported in mammalian cell lines such as HEK cells and teleost cell lines or live embryos such as zebrafish [10] [25] [34]. Here, we demonstrated that baculovirus can infect cloudy catshark primary cultured cells incubated at a low temperature with high osmotic pressure and urea concentration. This fact suggests that it may be possible to perform *in vivo* genetic manipulation using viral infection in cartilaginous fishes, and confirms that baculovirus can infect a wide range of hosts. The viral titer of primary cultured cells from cloudy catshark embryos was 1.36 × 10^5^ gtu/ml, which is comparable to that of HEK293A cells (1.55 × 10^5^ gtu/ml).

After adenovirus infection, on the other hand, we observed no GFP signal in cloudy catshark cells. Adenovirus is known to infect and induce gene expression in teleost cell lines *in vitro* and in zebrafish and medaka *in vivo* [11] [12]. This raises the possibility that infectivity of adenovirus is low in cartilaginous fish cells. Alternatively, the culture conditions with high osmolality, high urea, and low temperature might have reduced infectivity of adenovirus in the present study.

Electroporation and lipofection have been demonstrated to be effective not only in mammalian cell lines but also in teleost, invertebrate and dogfish shark cell lines [35] [36] [37] [38]. The present study shows that these methods can also be applied to cartilaginous fish primary cultured cells, which are suggested to retain more of the physiological properties of the original cells than conventional established cell lines do. In keeping with the results obtained by Parton *et al*., in which the transfection rate using electroporation or Cellfectamine^TM^, another commercially available lipofection reagent, into a dogfish shark cell line was 4% at best [8], the transfection rate to cloudy catshark cells in this study was equivalently low (Table 1) Meanwhile, PEI transfection, which is a method of chemical gene transfection similar to lipofection, failed to transfer an exogenous gene into cloudy catshark cells. Since PEI is thought to promote transfection through a similar process as lipofectamine, namely, by forming complexes with DNA and entering cells through endocytosis [39], further optimization may improve the efficacy of transfection by PEI.

### In vivo electroporation effectively introduced exogenous genes into cloudy catshark embryos and adult testis

We found that *in vivo* electroporation effectively introduces exogenous DNA fragments into cloudy catshark embryos. GFP was expressed strongly enough to be easily recognized through a fluorescent dissecting microscope. Some cloudy catshark embryos whose bodies were subjected to electroporation expressed GFP in the skeletal muscle cells (Fig. 5B); this expression was confirmed by EGFP immunohistochemistry of the transverse sections, indicating that the EGFP-expressing cells were restricted to the injected areas. We also successfully induced GFP expression in other parts including brain (Additional file 2). We could not validate the integration into the genome because long-term rearing was not possible in present culture method. Future improvements in culture methods or the use of inverse PCR may make it possible to determine the genomic integration of endogenous gene.

Previously, dyes such as DiI have been used as tracers in lineage tracing analyses in cartilaginous fishes [40] [41], which made it difficult to label specific cell types. In combination with the use of gene-specific promoters, electroporation enables cellular linage tracing of specific cells during development. Introduction of transgenes makes researchers possible to perform cell type specific labeling in living tissues by using enhancers/promoters, which has been used in many model species [42] [43] [44] [45] [46]. In spite of its effectiveness for transfection into culture cells *in vitro*, baculovirus failed to introduce a detectable level of GFP into tissues *in vivo*. The reason for this discrepancy remains to be determined, but the difference in circumstances between *in vitro* and *in vivo* situations, such as the immune system and intercellular adhesion *in vivo*, might be involved.

Furthermore, we succeeded in introducing a gene into the adult testis by electroporation through a small incision. Also, we succeeded in gene introduction in a part of the intestine and liver (Additional file 4), which suggests this method can be applied to various tissues, perhaps with some modification. Also, based on this introduction method of DNA fragments, functional analysis by gene overexpression and/or knockdown in the cells of adult tissues can be applied in the future.

On the other hand, in the present study, we could not find gene-introduced germ cells in the testis with electroporation although we succeeded in transducing EGFP-expression vector into cells outside spermatocysts. One possible reason for this is the presence of basement membrane lining the spermatocyst structure. Since it has been reported in other sharks that spermatocysts are lined with a basement membrane [1] [19], it is possible that the plasmid injected to the stroma did not penetrate into the spermatocysts. In other cartilaginous fishes such as spiny dogfish shark (*Squalus acanthias*) and bonnethead shark *(Sphyrna tiburo*), it has been reported that more undifferentiated germ cells (primary spermatogonia) do not have a cyst structure [1] [47] [48]. If such undifferentiated germ cells are identified in cloudy catshark testis in the future, gene introduction to germ cells by electroporation will become more feasible.

### Possible future applications of gene transfer methods in cartilaginous fishes

In the present study, we established a primary culture system for cloudy catshark cells and successfully introduced exogenous genes by means of lipofection, electroporation and baculovirus infection *in vitro*. These achievements will allow us to apply these gene transfer methods in various future studies investigating the development and physiology of cartilaginous fishes at the cellular level. For example, previous functional studies of the transporters or channels involved in osmotic regulation have been performed using *Xenopus* oocytes [49]. However, because the intracellular and extracellular environments of cartilaginous fishes are completely different from those of amphibians, it has thus far been difficult to accurately analyze the activities of these transporters and channels in cartilaginous fishes. The methods described here will enable us to analyze the activities of transporters and channels in more nearly physiological conditions. Although the efficiency of these transfection methods is still low, single-cell analysis methods such as patch clamp and Ca^2+^/cAMP imaging can be applied.

We also succeeded in *in vivo* gene transfection in both embryos and adults of cartilaginous fishes by means of electroporation, which allows *in vivo* forced expression locally at the site of electroporation. In the future, the introduction of a Cas9-gRNA integrated vector [50] may permit the generation of cloudy catsharks with knockout of a specific gene by the CRISPR/Cas9 system at local sites. Furthermore, future genetic modification in testicular germ cells could lead to the generation of genetically modified individuals. These techniques will likely use a reverse-genetics approach in cartilaginous fishes. Future improvement of transfection efficiency and the development of a means of transfection to germ cell lines will make generation of transgenic cloudy catsharks more realistic.

## Conclusions

We established a primary cell culture and performed exogenous gene introduction into embryos of a promising model cartilaginous fish, cloudy catshark, both *in vitro* and *in vivo*. Furthermore, we succeeded for the first time in transducing exogenous DNA constructs into adult cloudy catshark tissues. These methods can be applied to future physiological analyses at the cellular, tissue, and individual levels, enabling us to explore the molecular mechanisms involved in various phenomena that have yet to be clarified in cartilaginous fishes. Further, these methods should be the first step toward the production of transgenic cartilaginous fishes.

## Declarations

### Ethics approval and consent to participate

All experiments were conducted in accordance with the Guidelines for Care and Use of Animals approved by the ethics committee of the University of Tokyo (P19-2).

### Consent for publication

Not applicable

### Availability of data and materials

All data generated or analyzed during this study are included in this published article.

## Authors’ contributions

CF, SK and SH designed the research. CF, CU, MC and SK performed experiments. SI and HB contributed to the generation of baculovirus. SK and SH supervised the projects. CF prepared the draft of manuscript and SK, CU, SI and SH modified the manuscript. All authors read and approved the final manuscript.

## Acknowledgements

We are grateful to all the staff of Oarai Aquarium for their kind support in providing cloudy catshark embryos. We also thank Drs. Yuki Honda (Regional Fish Institute), Wataru Takagi (University of Tokyo), Mr. Naotaka Aburatani (University of Tokyo) and Mr. Koya Shimoyama (University of Tokyo) for helpful advice and assistance with the care and manipulation of cloudy catsharks. We also thank Drs. Yu Toyoda, Tappei Takada, Misaki Miyoshi, and Yuko Mochizuki (University of Tokyo) for their technical support and helpful advice regarding adenovirus production.

## Authors’ information

Chika Fujimori, Susumu Hyodo, Shinji Kanda:

Atmosphere and Ocean Research Institute, The University of Tokyo, Kashiwa, Chiba 277-8564, Japan

Chika Fujimori

Present address: Optics and Imaging Facility, National Institute for Basic Biology, Nishigonaka 38, Myodaiji, Okazaki 444-8585 Aichi, Japan

Chie Umatani:

Department of Biological Sciences, Graduate School of Science, The University of Tokyo, Bunkyo-ku, Tokyo 113-0033, Japan

Misaki Chimura, Shigeho Ijiri:

Graduate School of Fisheries Sciences, Hokkaido University, Minato-cho 3-1-1, Hakodate, Hokkaido 041-8611, Japan

Hisanori Bando:

Division of Applied Bioscience, Research Faculty of Agriculture, Hokkaido University, Sapporo 060-8589, Japan

## Corresponding authors

Chika Fujimori and Shinji Kanda

### Additional file 1

(avi file) Time-lapse images of primary cultured cells from cloudy catshark embryos. After seeding of dissociated cells into gelatin-coated wells, images were acquired every 5 mins for 25 hours. In these time-lapse recording, cell divisions were rarely observed. The movie runs at 15 frames (75 min) per second.

### Additional file 2

(pdf file) *In vivo* gene introduction into cloudy catshark embryos by electroporation. **A** In other cloudy catshark embryos whose brains had been subjected to electroporation, GFP fluorescence was observed at the injected site. **B** EGFP immunohistochemistry using transverse section of **A** indicated that EGFP-immunoreactive neurons were localized in the subpallium. (**C, D**) show magnified images of the boxed regions of (**B**). **E**, Fluorescence images of other embryos seven days after electroporation in the brain or nervous system. Note that **A** shows ventral view of the embryo head whereas **B-D** represent the transverse section of them. Scale bars represent 500 μm (**B**) and 20 μm (**C, D**).

### Additional file 3

(pdf file) Morphology of adult cloudy catshark testis. **A** Dorsal side image of adult cloudy catshark testis. Yellow arrows show germinal zone where plasmid was injected prior to the electroporation. **B** Hematoxylin and eosin staining of transverse sections of adult cloudy catshark testis. Because of the large size of the cloudy catshark testis, the testis was cut into two parts, medial (left) and lateral (right), and cryosections were prepared. Arrow represents germinal zone and magnified image of this area was shown in **C**. Scale bars represent 5 mm (**B**), 200 μm (**C**), respectively.

### Additional file 4

(pdf file) *In vivo* gene introduction into adult cloudy catshark tissues. Similar to *in vivo* electroporation to testis, liver or intestine was exposed from small incision in the abdomen and 50-100 μg of pEGFP-N1 vector was injected. Then, electroporation was applied at 30 V for 50 ms with 5 pulses using bipolar electrode. Fluorescence images of adult cloudy catshark intestine and liver seven days after electroporation were shown at **A** and **B**, respectively. **C, D** Fluorescence images of untreated intestine (**C**) and liver (**D**). Scale bars represent 1 mm.

