## Supplementary figures and images for "*In vitro* and *in vivo* gene introduction in the cloudy catshark (*Scyliorhinus torazame*), a cartilaginous fish"

### Additional file 2

Additional file 2.

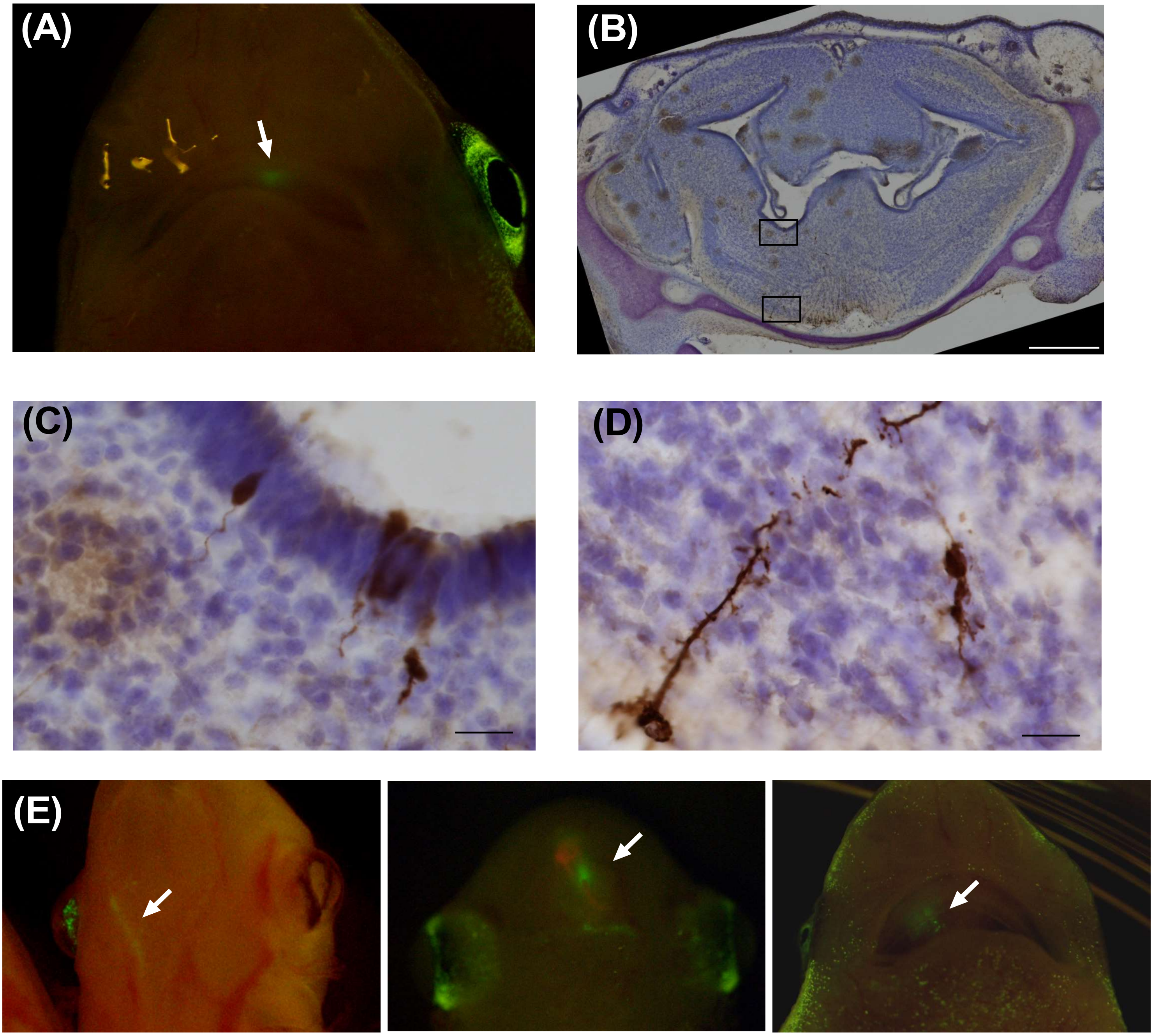

### Additional file 3

Additional file 3

(A)

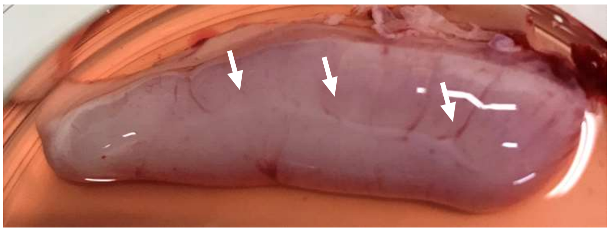

(B)

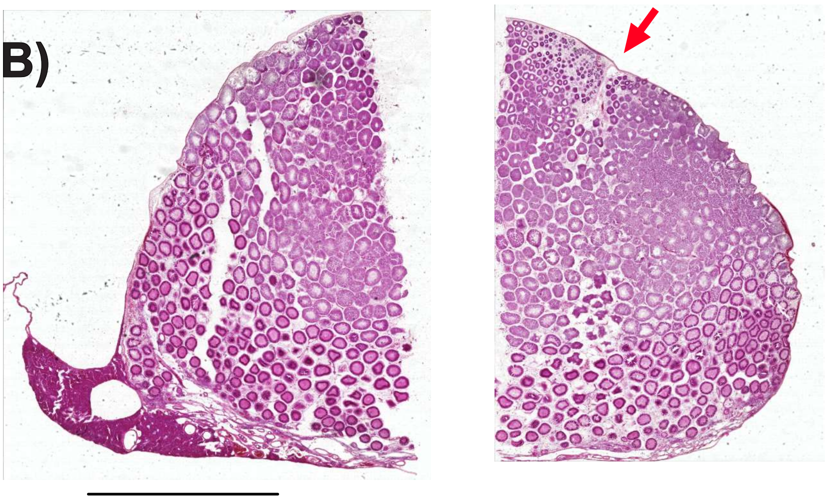

(C)

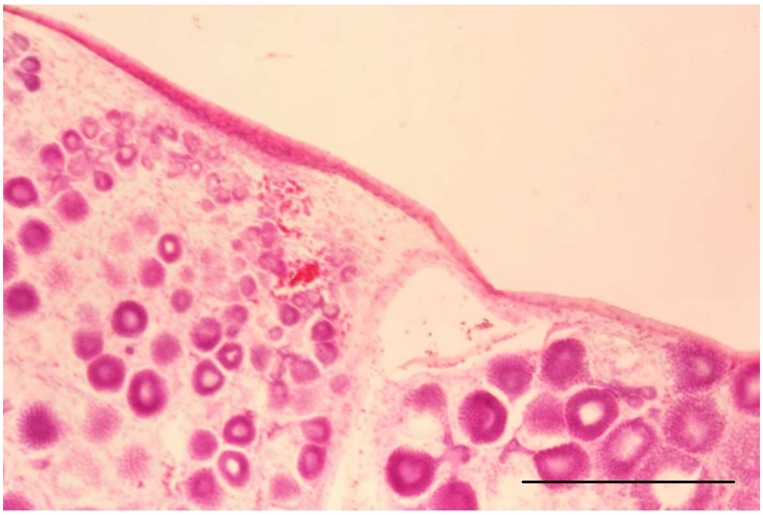

### Additional file 4

Additional file 4

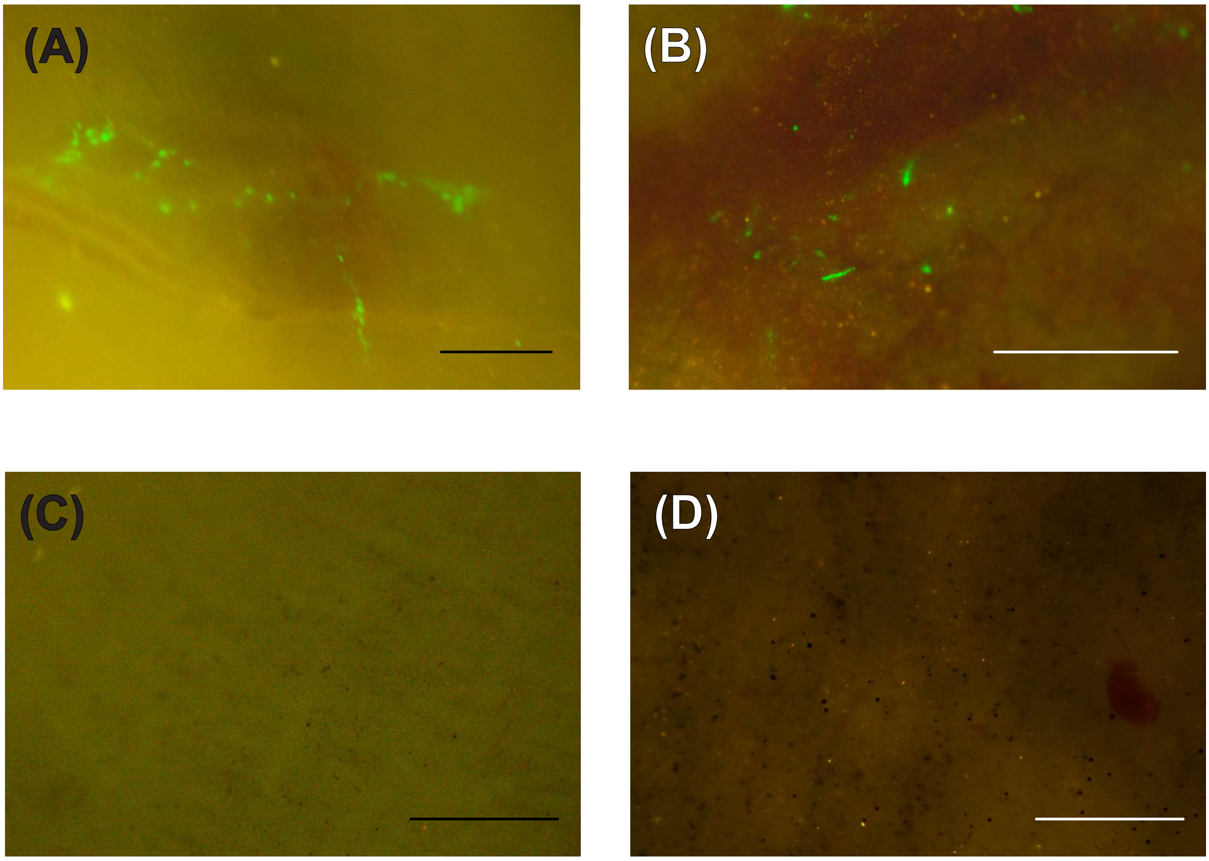
